# Elucidation of sub-cellular H_2_S metabolism in *Solanum lycopersicum* L. and its assessment under development and biotic stress

**DOI:** 10.1101/2021.09.25.461755

**Authors:** Aksar Ali Chowdhary, Sonal Mishra, Vikram Singh, Vikas Srivastava

## Abstract

The signalling molecules serve as a fundamental requirement in plants and respond to various internal and external cues. Among several signalling molecules, the significance of gasotransmitters has been realized in several plant developmental and environmental constraints. The hydrogen sulfide (H_2_S) is a novel signalling molecule in higher plants and is involved in several physiological processes right from seed germination to flowering and fruit ripening. Moreover, H_2_S also assist plants in managing biotic and abiotic stresses, therefore serves as one of the imperative choice of chemical priming. Yet, the metabolism of H_2_S is not much explored and only appraisal study is made till date from *Arabidopsis thaliana*. Therefore, the present investigation explored the elucidation of H_2_S metabolism in crop plant *Solanum lycopersicum* L. Through in silico investigations the study demonstrated the participation of 29 proteins involved in H_2_S metabolism, which are mainly localized in cytosol, chloroplast, and mitochondria. Additionally, the relevant protein-protein interactomes were also inferred for sub-cellular compartments and expression data were explored under development and biotic stresses namely PAMPs treatment and bacterial infection. The information generated here will be of high relevance to better target the H_2_S metabolism to enhance the tomato prospects and also serve a preliminary investigation to be adopted in other agronomic important crops.

## 1. Introduction

The assessment of hydrogen sulfide (H_2_S - a gasotransmitter; with characteristic rotten egg like smell) in life forms has dramatically changed from a pollutant (toxic) to an intermediate during sulfur assimilation into cysteine, and has also proven currently as a signalling component (Filipovic and Jovanovic, 2017). Like other gasotransmitters (NO and CO), H_2_S is also known to be able to cross cell membrane without receptors (Guo et al., 2016). The functionality of H_2_S primarily depends on its concentration and offer toxicity and signalling role at high and low concentrations, respectively (Dooley et al., 2013). The function of H_2_S-mediated signaling is a widespread incident and is conserved throughout diverse life forms. In animals, H_2_S was reported to function as neuro-modulator in brain cells and found to be involved in physiological and pathological processes, such as apoptosis and inflammatory progression (Jin and Pei et al, 2015). In plants, the contribution of H_2_S is observed in the regulation of several developmental traits and protection against several abiotic and biotic stresses (Corpas, 2019). The major physiological and developmental process in which H_2_S regulation was observed includes seed germination, organogenesis, stomatal closure/aperture, modulation of photosynthesis, autophagy regulation etc (Aroca et al., 2018; Arif et al., 2021). The H_2_S–mediated protection and tolerance was observed in almost all category of abiotic stress including oxidative (Corpas, 2019; Corpas and Palma, 2020), heavy metal (Li et al., 2012; Kharbech et al., 2017; Kharbech et al., 2020), salt (Mostofa et al., 2015; Ding et al., 2019), drought (Chen et al., 2016; Zhou et al., 2020), heat (Zhou et al., 2018), chilling (Pan et al., 2020) and waterlogging (Xiao et al., 2020) stress. Apart, the contribution of H_2_S was also observed for the development of resistance against phytopathogens (Vojtovič et al., 2020). Due to its immense potential for the development of stress tolerance features, like phytohormones (Mishra et al., 2020) and GABA (Srivastava et al., 2021) it is also useful for chemical priming to develop cross-adaption for stress tolerance (Ahmed et al., 2021). Besides, its contribution towards enhanced production of metabolites was also reported (Li et al., 2016). Additionally, the H_2_S also trigger post translation modifications (through persulfidation), which lead to the significant modulation in the activities of proteins/enzymes being persulfidated (Park et al., 2015; Filipovic et al., 2018). Further a study reported that under baseline condition, 5% of Arabidopsis proteome may undergo persulfidation (Aroca et al., 2015; Aroca et al., 2017). Such persulfidation may lead to either activation (APX, LCD, RBOHD etc.) or inhibition (Catalase, NADP-Isocitrate dehydrogenase etc.) of concerned enzymes (Aroca et al., 2015; Corpas et al., 2019; Shen et al., 2020; Munoz-Vargas et al., 2018 & 2020). These studies hence suggested that the H_2_S offer their biological potential by regulation of ROS and antioxidant machinery, which are the important signalling component for plant development and stress response (Mishra et al., 2017). The sub-cellular locations of H_2_S metabolism was primarily reported to be in the cytosol, chloroplast and mitochondria (Gonzalez-Gordo et al., 2020), and involve several enzymatic components for H_2_S production and consumption (Li, 2015; Gonzalez-Gordo et al., 2020; Arif et al., 2021).

Though several aspects of H_2_S mediated stress tolerance and individual H_2_S enzyme associated biological features are known in plants, yet the cumulative account of the enzymatic sub-cellular network for its metabolism is poorly established, except for the Arabidopsis. (Gonz’alez-Gordo et al., 2020). We here attempted to elucidate the H_2_S metabolism in *Solanum lycopersicum* L. (tomato, formerly *Lycopersicon esculentum* Mill.) and established their functional relevance. In the present study, the focus involves the identification of tomato H_2_S metabolism associated enzymes and their sub-cellular locations, towards developing a working model for H_2_S metabolism in tomato. Along, the compartment specific interactions of enzymes associated with H_2_S metabolism were also undertaken. Besides, the expression analysis of genes coding for proteins involved in H_2_S production and consumption have also been assessed under developmental and pathogenic situations using gene expression analysis. The study will offer ways to better understand the H_2_S metabolism in tomato and offer reasonable insights for the betterment of tomato productivity.

## 2. Materials and Methods

### 2.1. Database Search

National Center for Biotechnology Information (https://www.ncbi.nlm.nih.gov/) and Sol Genomic Network (https://solgenomics.net/) were used to retrieve the proteins associated with H_2_S metabolism in *S. lycopersicum* by using BLAST (BLASTP and TBLASTN) analysis. All the 26 proteins associated with H_2_S metabolism reported in *A. thaliana* (Gonz’alez-Gordo et al., 2020) were used as query. The retrieved proteins of tomato with a highly stringent E-value, namely, less than 10^−10^ was considered for the study. Further, these proteins were also subjected to the reverse BLAST in TAIR (https://www.arabidopsis.org/Blast/index.jsp), to identify their closest orthologs in Arabidopsis. All the retrieved sequences were then subjected for the functional domain analysis using NCBI-CDD search, to filter out any proteins not related to H_2_S metabolism. The identified proteins were named either (i) as reported earlier in tomato, and/or (ii) considering the information as given in the Arabidopsis model (Gonz’alez-Gordo et al., 2020; Hu et al., 2020; Liu et al., 2019). The functional assignment of the identified proteins was done by considering information from solgenomics (https://solcyc.solgenomics.net/), orthologs function in Arabidopsis, CDD search and BLAST results. All the identified proteins were given names according to their appearance in the genome; however, the previously assigned names were kept as such (Liu et al., 2019; Hu et al., 2020).

### 2.2. Sub-cellular localization

The sub-cellular localization of proteins was predicted based on their analysis using 12 independent sub-cellular localization tools viz, WoLF PSORT (Horton et al., 2007), Deeploc 1.0 (Armenteros et al., 2017), ChloroP 1.1. (Emanuelsson et al., 2019), TargetP 2.0 (Armenteros et al., 2019a), Plant-mSubP (Sahu et al., 2020), BUSCA (Savojardo et al., 2018), LOC TREE 3 (Goldberg et al., 2014), DeepMito (Savojardo et al., 2020), LOCALIZER (Sperschneider et al., 2017), CELLO2GO (Yu et al., 2014), SignalP-5.0 (Armenteros et al., 2019b) and DeepSig (Savojardo et al., 2018). Out of them, most of the predictions from SignalP-5.0 and DeepSig, were falling into “other” categories, thus not considered further. The information from remaining 10 prediction tools were used to develop a confidence score (Number of occurrence at particular location/ Total number of prediction tools explored). A particular protein was assigned the sub-cellular location with maximum confidence score. However, for the proteins predicted to be localized in two sub-cellular compartments, support from sub-cellular localization of their orthologous proteins in Arabidopis and information from previous research studies were also taken into consideration (Liu et al., 2019; Hu et al., 2020).

### 2.3. PPI Network Analysis

To further substantiate the functional pertinence of *S. lycopersicum* proteins under investigation, conserved domains were searched in these proteins identified using NCBI Conserved Domain Search-NIH Database (https://www.ncbi.nlm.nih.gov/Structure/cdd/wrpsb.cgi). The sequences were then considered for PPI (protein-protein interaction) network analysis. The STRING database v11 (Szklarczyk et al., 2019) was used for the analysis of protein–protein interaction, using compartment specific identified proteins as inputs for analysis. The confidence view was generated by setting the filter to medium confidence (0.400).

### 2.4. Gene Expression Analysis

The expression analysis of genes associated with the production and consumption of H_2_S (between cysteine and H_2_S) was performed using RNA-Seq data, available in tomato functional genomic database (http://ted.bti.cornell.edu/cgi-bin/TFGD/digital/home.cgi). The assessment of genes associated with cytosolic H_2_S metabolism utilizes the expression of *SlLCD1, SlOAS1, SlOAS2, SlOAS3, SlOAS4, SlLCD2, SlOAS6* and *SlOAS9*. The expression data of *SlDCD2, SlNFS2, SlOAS5* and *SlSiR1* were used for the study of chloroplastic H_2_S metabolism, and of *SlDCD1, SlOAS7* and *SlNFS1* were analyzed for mitochondrial H_2_S metabolism. For the assessment of tissue specific expression, data from D004 experiment was utilized. In brief, the Heinz cultivar was used for the assessment of the expression of associated genes under different development stages viz, unopened flower buds, fully opened flowers, 1 cm fruits, 2 cm fruits, 3 cm fruits, mature green fruits, breaker fruits, breaker fruits+10 fruits, leaves and roots. Rio Grande prf3 (deletion in Prf) cultivar was used to assess the response under PAMP/biotic stress (D007 experiment). In all the experiments mock samples comprised of MgCl_2_. The expression analysis during PAMPs includes two durations (30 min and 6h) and involve treatments of leaves with cold shock protein 22 (csp22), flagellin 22 (flg22), flagellin II-28 (flgII-28), Lipopolysaccharides (LPS) and Peptidoglycan (PGN). The bacterial treatment includes the leaf infection with *Pseudomonas syringae* (DC3000), *P. fluorosence, P. putida* and *Agrobacterium tumifaciens*. The expression data was assessed at 30 min and 6 h (Rosli et al. 2013). The expression data was visualized using heat map developed by MeV (Howe et al., 2010).

## 3. Results

### 3.1. Identification of H_2_S metabolism related candidate proteins in tomato

During cysteine biosynthesis H_2_S serve as an intermediate, and over the time its significance in signalling has been observed in many aspects of plant biology (Corpas, 2019; Liu et al., 2021). The metabolism of H_2_S is not established properly in plants; however Gonzalez-Gordo et al., (2020) recently attempted to identify proteins involved in H_2_S metabolism in *A. thaliana* and presented the working model utilizing proteins directly or indirectly related to the cysteine metabolism. We here attempted to bring together the information of H_2_S metabolism related proteins in *S. lycopersicum* and identification of the remaining candidate proteins through in silico investigation utilizing information of *A. thaliana* proteins. The contribution of cytosol in the H_2_S metabolism is well known. In the tomato cytosol, the H_2_S metabolism (Table 1) involves the participation of L-Cysteine desulfhydrase (SlLCD1 and SlLCD2), Cysteine synthase / O-acetyl-l-serine (thiol)lyase (SlOAS1, SlOAS2, SlOAS3, SlOAS4, SlOAS6 and SlOAS9), and Selenium binding protein (SlSBP1).

**Table 1.**
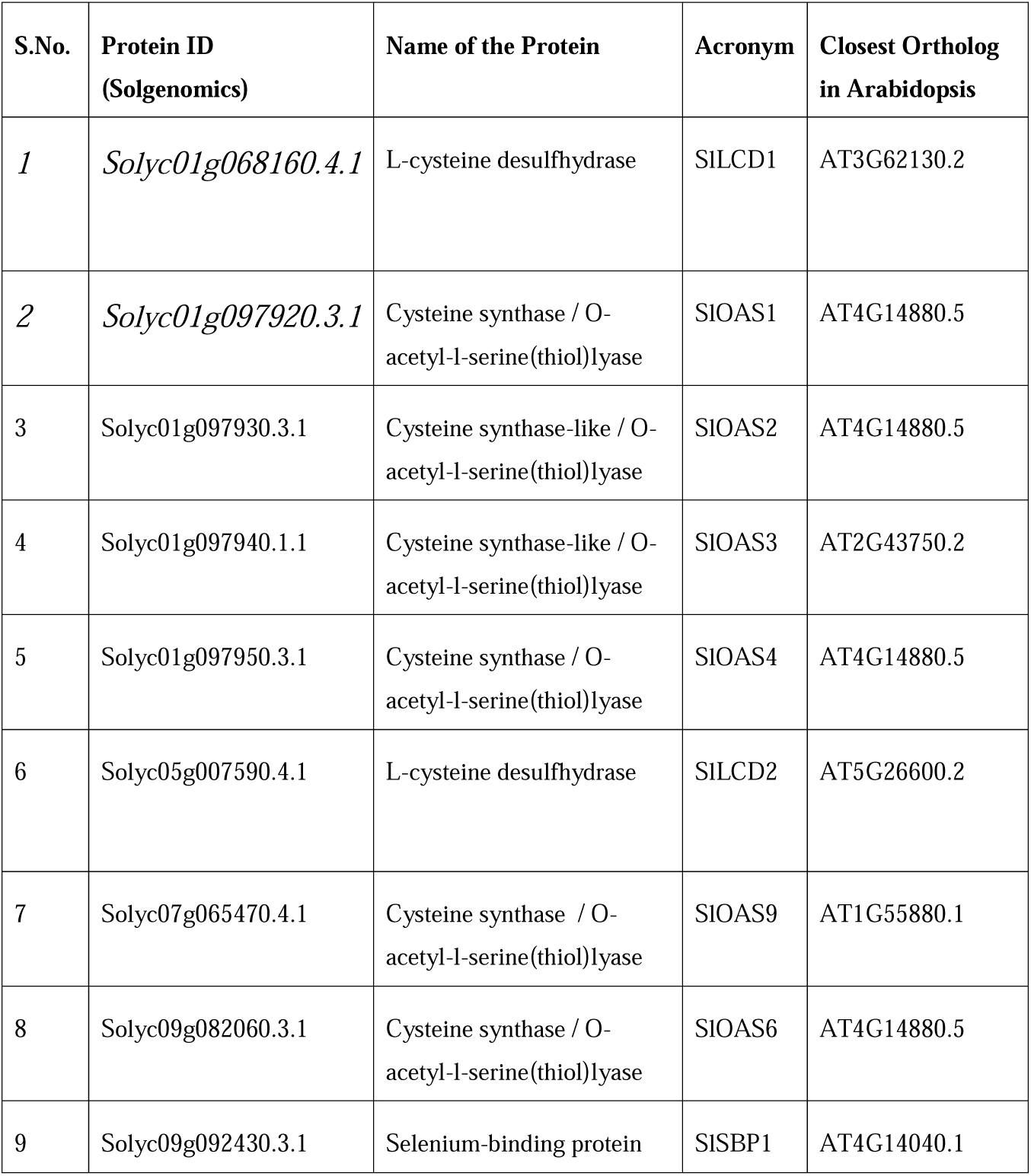
Cytosolic enzymes involved in H_2_S metabolism in *S. lycopersicum*.

The chloroplast is the one of the most important sites of sulfur assimilation into amino acids, which involves sulfate reduction and cysteine production (Birke et al., 2015). Table 2 illustrates the identified enzymes of tomato involved in chloroplastic H_2_S metabolism. In total, 12 enzymes were identified which includes Cystathionine gamma synthase (SlCGS1), Thiosulfate sulfurtransferase (SlMST1, SlMST2 and SLMST4), D-cysteine desulfhydrase (SlDCD2), Cysteine desulfurase (SlNFS2), Lipoyl synthase (SlLIP1 and SlLIP3), Cysteine synthase / O-acetyl-l-serine(thiol)lyase (SlOAS5), Cystathionine beta-lyase (SlCBL2 and SlCBL3) and Sulfite reductase (SlSiR1). The contribution of H_2_S is evident in mitochondria as it affects mitochondrial electron transport chain by inhibiting cytochrome c oxidase (Complex IV). It contributed significantly in processes related to energy production and cellular ageing, particularly during drought stress (Birke et al., 2013; Jing et al., 2018). Our analysis suggested six enzymes responsible for mitochondrial H_2_S metabolism, viz. D-cysteine desulfhydrase (SlDCD1), Cysteine synthase / O-acetyl-l-serine (thiol) lyase (SlOAS7), Cysteine desulfurase (SlNFS1), Bifunctional L-3-cyanoalanine synthase/ cysteine synthase (Solyc04g058120.3.1, SlCYSC2; not considered further as its expression was not observed in any of the developmental tissue), Lipoyl synthase (SlLIP2) and Bifunctional L-3-cyanoalanine synthase/ cysteine synthase / O-acetyl-l-serine (thiol) lyase (SlOAS8) (Table 3). The location of Solyc06g009860.1.1 (named as SlMST3) was observed as extracellular, which possess mercaptopyruvate sulfurtransferase activity. Besides, we also could not get the localization consensus for Solyc04g055230.2.1, which appears to be Cystathionine-β-lyase (named as SlCBL1). Both of these proteins (SlMST3 and SlCBL1) were not used for the current model, though their functionality in tomato cannot be denied. The proteins localized in cytosol, chloroplast and mitochondria were utilized for the depiction of putative working model of H_2_S metabolism in tomato (Fig. 1).

**Table 2.**
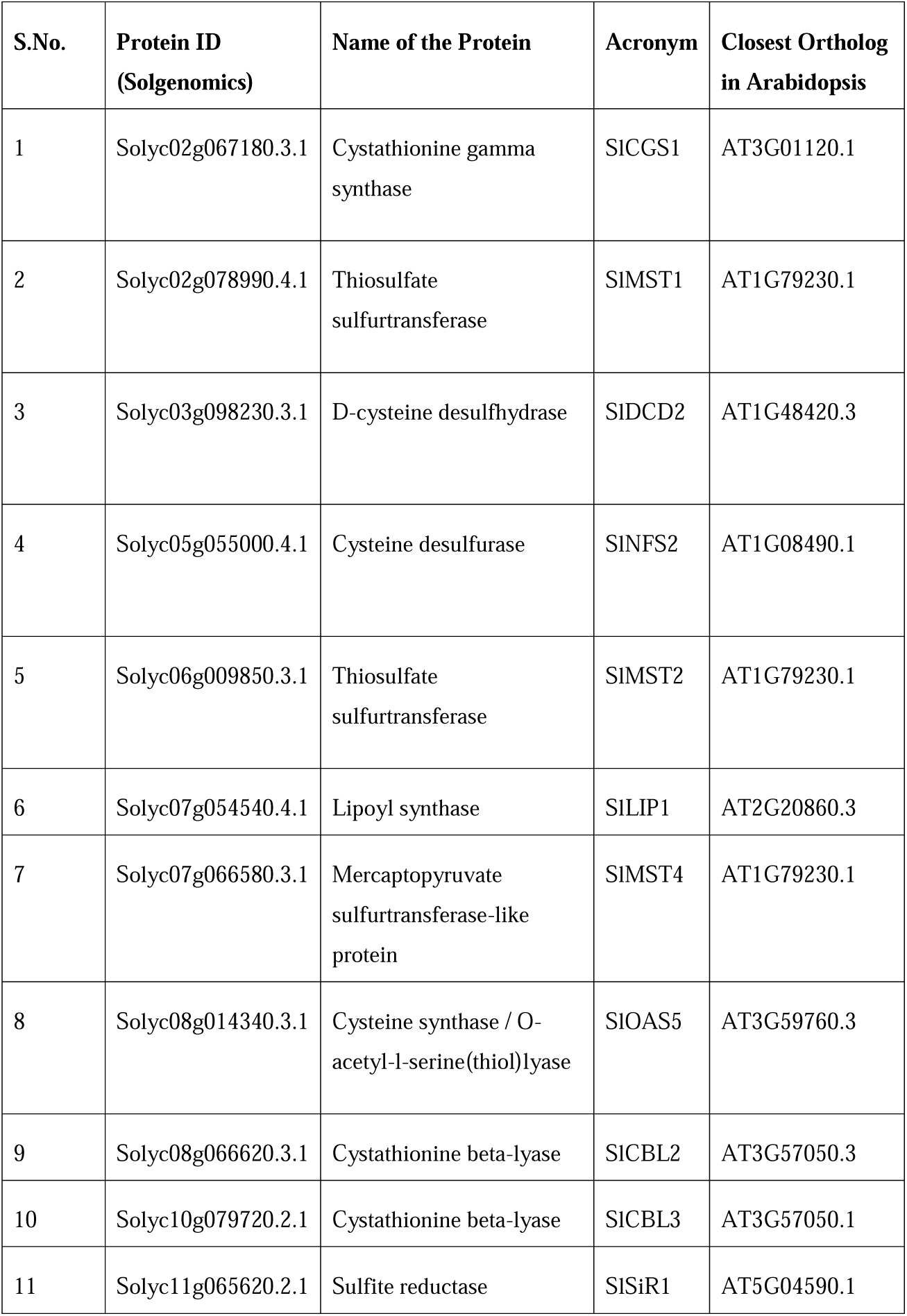
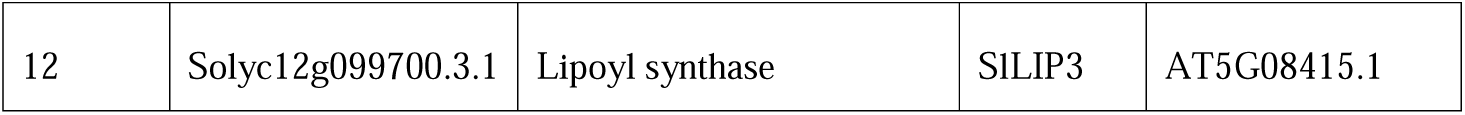
Chloroplastic enzymes involved in H_2_S metabolism in *S. lycopersicum*.

| S.No. | Protein ID<br>(Solgenomics) | Name of the Protein | Acronym | Closest Ortholog<br>in Arabidopsis |
| --- | --- | --- | --- | --- |
| 1 | Solyc02g067180.3.1 | Cystathionine gamma<br>synthase | SICGS1 | AT3G01120.1 |
| 2 | Solyc02g078990.4.1 | Thiosulfate<br>sulfurtransferase | SIMST1 | AT1G79230.1 |
| 3 | Solyc03g098230.3.1 | D-cysteine desulphydrase | SIDCD2 | AT1G48420.3 |
| 4 | Solyc05g055000.4.1 | Cysteine desulfurase | SINFS2 | AT1G08490.1 |
| 5 | Solyc06g009850.3.1 | Thiosulfate<br>sulfurtransferase | SIMST2 | AT1G79230.1 |
| 6 | Solyc07g054540.4.1 | Lipoyl synthase | SILIP1 | AT2G20860.3 |
| 7 | Solyc07g066580.3.1 | Mercaptopyruvate<br>sulfurtransferase-like<br>protein | SIMST4 | AT1G79230.1 |
| 8 | Solyc08g014340.3.1 | Cysteine synthase / O-acetyl-l-serine(thiol)lyase | SIOAS5 | AT3G59760.3 |
| 9 | Solyc08g066620.3.1 | Cystathionine beta-lyase | SICBL2 | AT3G57050.3 |
| 10 | Solyc10g079720.2.1 | Cystathionine beta-lyase | SICBL3 | AT3G57050.1 |
| 11 | Solyc11g065620.2.1 | Sulfite reductase | SISiR1 | AT5G04590.1 |
| 12 | Solyc12g099700.3.1 | Lipoyl synthase | SILIP3 | AT5G08415.1 |

**Table 3.** Mitochondrial enzymes involved in H_2_S metabolism in *S. lycopersicum*.

| S.No. | Protein ID<br>(Solgenomics) | Name of the Protein | Acronym | Closest<br>Ortholog in<br>Arabidopsis |
| --- | --- | --- | --- | --- |
| 1 | Solyc01g008900.4.1 | D-cysteine desulhydrase | SIDCD1 | AT3G26115.1 |
| 2 | Solyc01g094790.3.1 | Cysteine synthase / O-acetyl-l-serine(thiol)lyase | SIOAS7 | AT3G61440.1 |
| 3 | Solyc02g091900.4.1 | Cysteine desulfurase | SINFS1 | AT5G65720.2 |
| 4 | Solyc08g068440.4.1 | Lipoyl synthase | SILIP2 | AT2G20860.3 |
| 5 | Solyc10g012370.3.1 | Bifunctional L-3-cyanoalanine synthase/cysteine synthase / O-acetyl-l-serine(thiol)lyase | SIOAS8 | AT3G61440.1 |

**Figure. 1.**
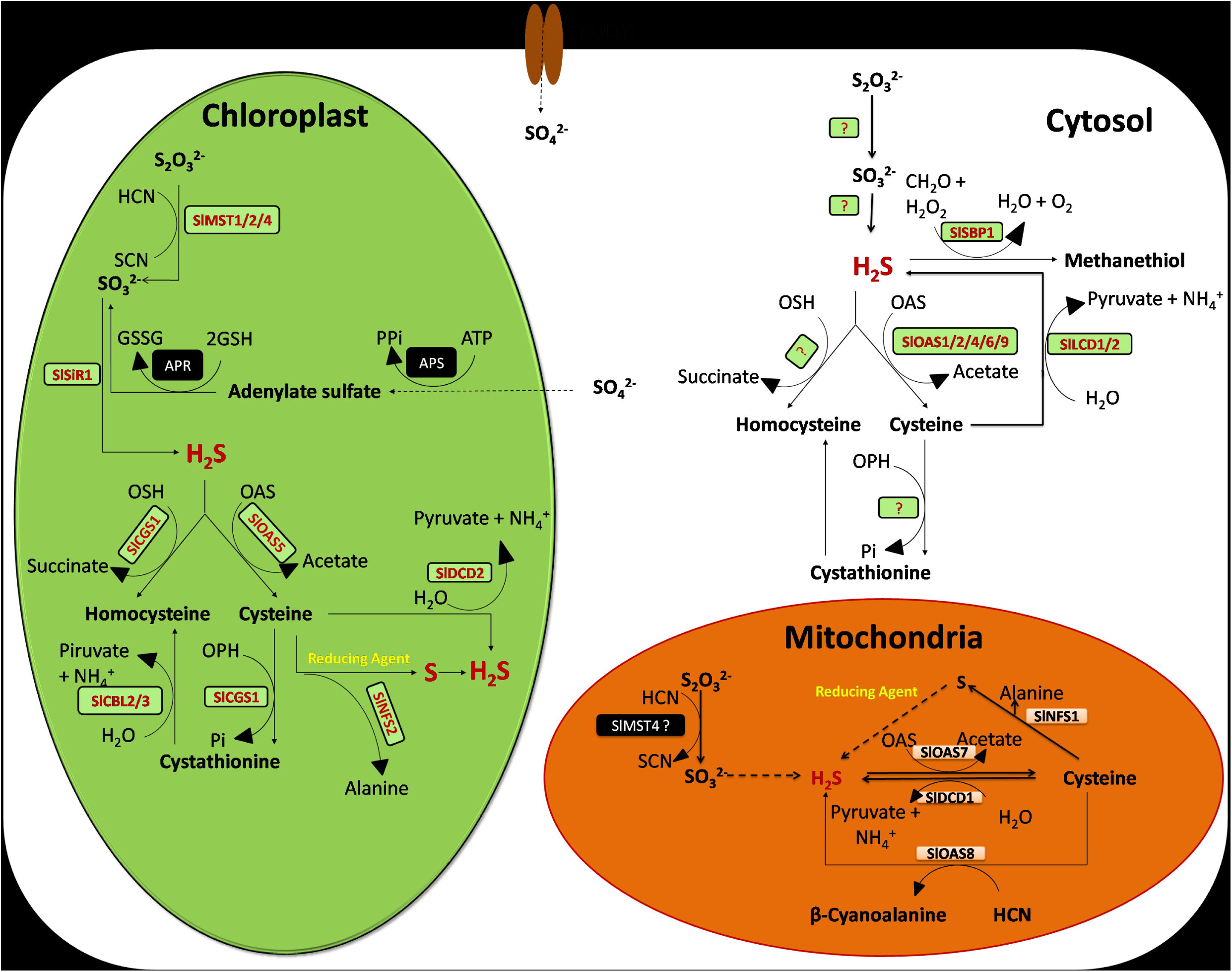
Working model for H_2_S metabolism in *S. lycopersicum*. Enzymatic reactions involved in H_2_S metabolism via cysteine (Cys) metabolism at different subcellular localization including cytosol, chloroplast, and mitochondrion in *S. lycopersicum*. SULTR, sulfate transporters; APS, ATP sulfurylase; APR, adenylylsulfate reductase. The enzymes in green and red box are identified in present study. The reactions need further confirmation is marked as question. Refer tables 1 and 2 and 3 to find the meaning of each abbreviation.

### 3.2. PPI of H_2_S metabolism related proteins in tomato

The sub-cellular protein-protein interaction (PPI) network was developed for the proteins localized in cytosol, chloroplast and mitochondria (Fig. 2). The network of cytosolic enzymes associated with H_2_S metabolism in *S. lycopersicum* demonstrated that out of nine proteins, eight proteins (except SlSBP1) i.e SlLCD1, SlOAS2, SlOAS6, SlOAS9, SlOAS1, SlLCD2, SlOAS3 and SlOAS4 exhibited a strong and direct interactions among themselves. However, no interaction was observed with SlSBP1, which corroborated with the findings in Arabidopsis (Gonzalez-Gordo et al., 2020). PPI network of chloroplast enzymes suggested that all the twelve proteins are interacted with each other, there by forming a network. However, some of the proteins have a strong interaction, namely, SlCBL3, SlOAS5, SlCGS1, SlMST1, SlMST4, SlDCD2, SlNFS2, SlSiR1, SlCBL2 while proteins SlLIP3, SlSiR1, SlDCD2 and SlLIP1 showed weak interactions with each other. Analysis of mitochondrial proteins interaction demonstrated that all the five proteins demonstrated interaction, but forming two groups. Further, strong interaction was observed between SlOAS7 and SlOAS8. The SlNFS1 exhibited interaction with SlLIP2 and SlDCD1, in which the former is comparatively stronger. The reasonable interactions among the enzymes involved in H_2_S metabolism corroborated with the findings in Arabidopsis (Gonzalez-Gordo et al., 2020).

**Figure. 2.**
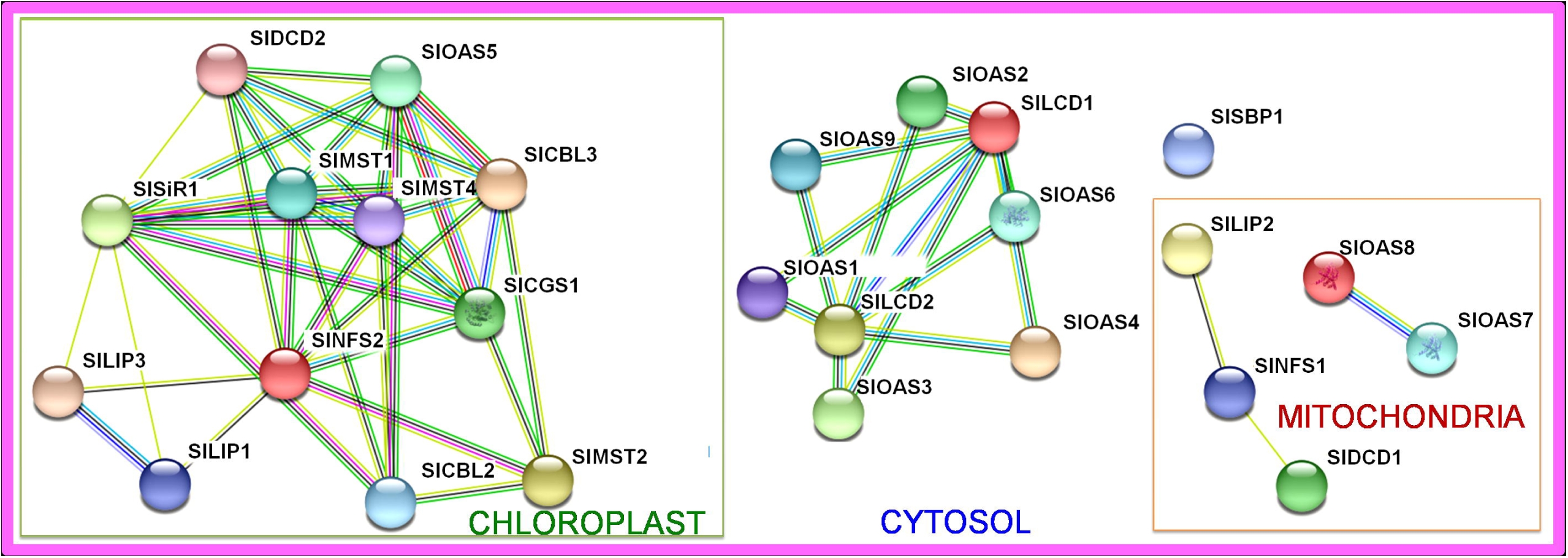
Protein-protein interaction (PPI) network of components involved in H_2_S metabolism in *S. lycopersicum*. Network nodes (circles) represent proteins and colored lines represent the types of interaction evidences used to predict the associations viz green line (indicates neighborhood evidence), blue line (indicates co-occurrence evidence), pink line (indicates experimental evidence), dark yellow line (indicates text-mining evidence), light blue line (indicates database evidence), black line (indicates co-expression evidence), purple line (indicates protein homology evidence). PPI Networks were done with STRING v11 (Szklarczyk et al., 2019). Refer tables 1 and 2 and 3 to find the meaning of each acronym.

### 3.3. Expression of genes associated with H_2_S metabolism

The in silico expression data available for Heintz and Rio Grande prf3 (deletion in Prf) cultivar was used for expression study under development and biotic stress, respectively. In particular, the focus here is to evaluate the expression of genes associated with either H_2_S production or consumption, to identify the significance of genes from each category under both conditions (Fig 3).

**Figure. 3.**
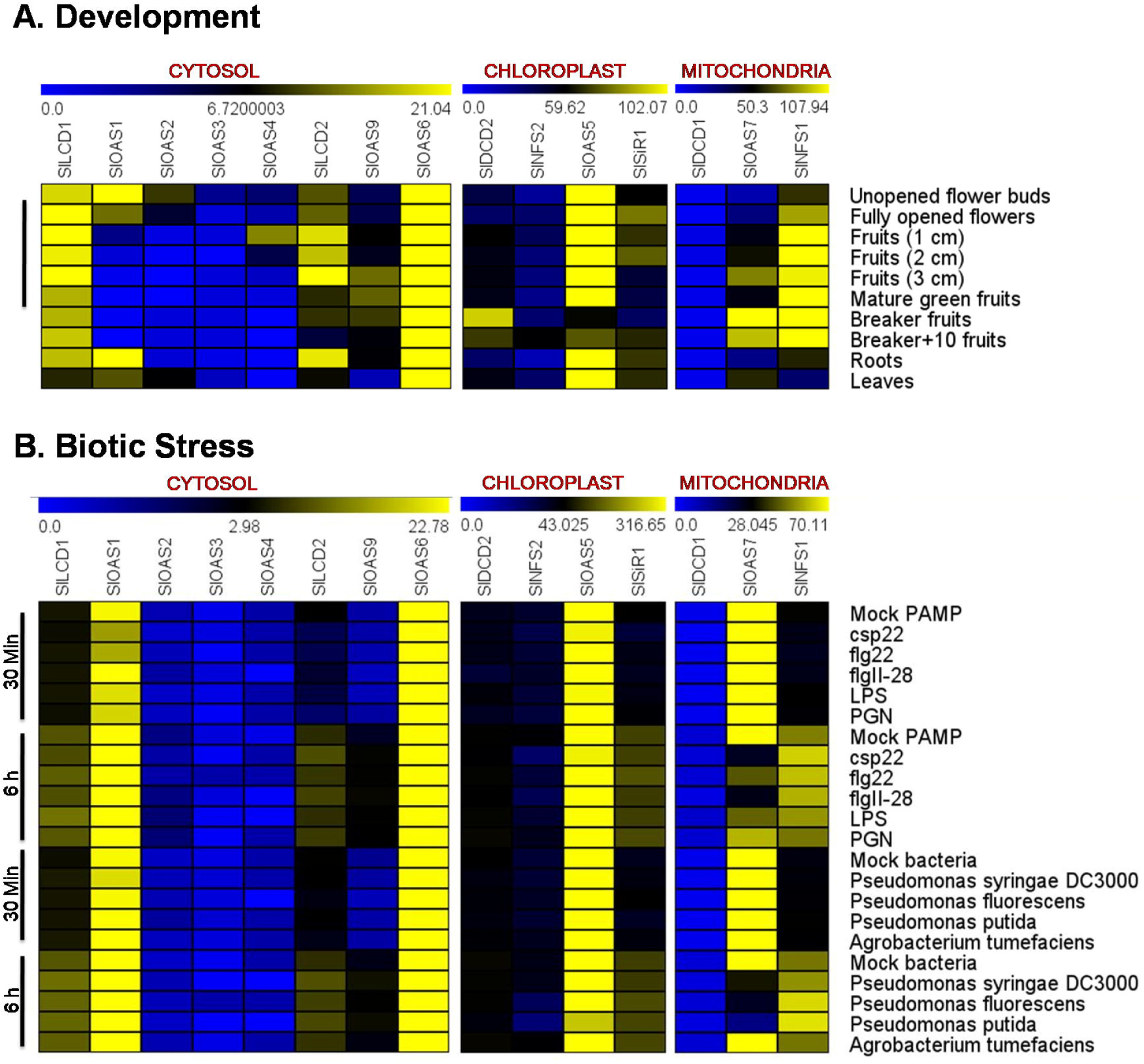
Heatmap representation for in silico expression. Relative expression analysis of the genes encoding enzymes related to H_2_S production and consumption, localized in cytosol, chloroplast and mitochondria of *S. lycopersicum*: (A) Development (B) Biotic stress-PAMP and bacterial treatment. Expression was visualized using heat maps by MeV (Howe et al., 2010). Blocks with colors indicate decreased (blue) or increased (yellow) transcript accumulation relative to the respective control. The bar at the top of the heat map represents relative expression values. The numbers below the bar indicate the log-transformed fold change. Refer tables 1 and 2 and 3 to find the meaning of each abbreviation.

#### 3.3.1. Expression under different developmental stages

All the genes exhibited differential expression at diverse developmental stages (Fig 3A). Among genes associated with cytosolic H_2_S metabolism, the highest expression of *SlOAS6* was observed in leaves. The expression of *SlLCD1, SlLCD2*, and *SlOAS9* demonstrated highest expression in ‘3 cm fruits’. The *SlOAS1/ SlOAS2/ SlOAS3* and *SlOAS4* demonstrated highest expression in ‘unopened flower buds’ and ‘1 cm fruits’, respectively. The expression of genes associated with chloroplast metabolism exhibited highest expression as follows: *SlOAS5* (leaves), *SlDCD2* (breaker fruits), *SlSiR1* (fully opened flowers) and *SlNFS2* (breaker+10 fruits). The genes associated with mitochondrial H_2_S metabolism exhibited highest expression in tissues such as *SlNFS1* (breaker+10 fruits), *SlOAS7* (breaker fruits), and *SlDCD1* (1cm fruits). The study suggested that H_2_S metabolism plays significant contribution in tomato developmental stages and is specifically involves in stages of fruit ripening.

#### 3.3.2. Expression in response to PAMP treatments

The expression analysis suggested that among genes associated with cysteine synthase activity, the contribution of *SlOAS6* is very high. During 30 min of PAMP treatment its expression was down-regulated, however, at late stage (6 h) it was observed significantly up-regulated. Amongst, the highest expression was observed in response to csp22 and lowest in response to PGN. Similar trend was observed in case of *SlOAS9*, and the highest expression was observed in response to flgII-28. In the *SlOAS1, SlOAS2* and *SlOAS3*, modulation was not much significant; however *SlOAS2* demonstrated up-regulation at flg28 and flg28, LPS & PGN at 30 min and 6 h, respectively. We could not get rpkm value for *SlOAS4* at 30 min; however, at 6 h this gene demonstrated down-regulation. The *SlLCD1* was also not much modulated at 30 min, however exhibited up-regulation at 6 h with few PAMP such as LPS. Among genes associated with chloroplast H_2_S metabolism, the *SlOAS5* and *SlNFS2* did not showed much deviation at 30 min; however, they exhibited slight down-regulation with all PAMPs at late stage (6h). The *SlSiR1* and *SlDCD2* exhibited up-regulation, with few PAMPs at 6 h. In mitochondria, the *SlOAS7* significantly demonstrated up-regulation and down-regulation during early (30 min) and late (6 h) stages, respectively which were reversed in *SlNFS1*. No significant modulation was observed for *SlDCD1* (Fig 3B).

#### 3.3.3. Expression in response to bacterial treatments

In cytosol, *SlOAS6* demonstrated modulations at 6h only and significant up-regulation was observed in the samples treated with *P. fluorescence* and *P. putida. SlOAS2* and *SlOAS9* also demonstrated up-regulation with bacterial treatments at both durations and only 6 h, respectively. *SlOAS3* demonstrated up-regulation at 30 min however at 6 h it was down-regulated for *Pseudomonas* sps and upregulated for *A. tumifaciens*. Not much modulation was observed for *SlOAS1* and *SlLCD1*. With different bacterial treatments, not much effect was noticed at 30 min for chloroplast gene *SlOAS5*; however at 6 h significant down-regulation was observed in samples treated with *Pseudomonas* sps, and up-regulation with *Agrobacterium tumifaciens*. Not much modulation was observed in case of *SlDCD2* and *SlSiR1*. In case of *SlNFS2*, down-regulation was noticed with *P. fluorescence* and *P. putida*. In mitochondria, the expression of *SlOAS7* suggested down-regulation for all *Pseudomonas* sps, and up-regulation for *A. tumifaciens* at 6 h. Further, *SlNFS1* demonstrated up-regulation mainly for *Pseudomonas* sps. In case of *SlDCD1* not much modulation was observed at 30 min; however at 6 h slight downregulation was observed with *P. syringae* and *P. putida* (Fig 3B).

## 4. Discussion

Among plant biomolecules, Sulfur (S) is the basic constituents of certain amino acids viz, cysteine, homocysteine and methionine, and some secondary sulfur molecules like glucosylates, phytochelatins, polysulfides and sulfolipids (Beinert, 2000; Münchberg et al., 2007; Queval et al., 2009; Shimojima, 2011; Takahashi et al., 2011; Romero et al., 2014; Fuentes-Lara et al., 2019). Further, under stress conditions, it also plays a significant contribution in ROS and RNS related metabolism with a tripeptide like glutathione (GSH) or S-nitrosoglutathione (GSNO) (Noctor et al., 2012; Corpas et al., 2013; Pivato et al., 2014; Hasanuzzaman et al., 2017). In plants, S is uptaken as sulfate (SO4^2-^) from soil through specific transporters (SULTR). At organ and tissue level, the localization of SULTR was reported in the roots and vascular tissues, respectively (Gigolashvili and Kopriva, 2014; Yamaguchi et al., 2020). At sub-cellular level, the SULTR in Arabidopsis was also reported from plastid, suggested its contribution in assimilation of SO4^2-^, along with cytosol. However, other cellular component mitochondria and peroxisome also participated in S metabolism (Gonzalez-Gordo et al., 2020). The assimilation of SO4^2-^ in S metabolism involves the contribution of cytosol, chloroplast, mitochondria and peroxisome; utilizing inorganic S forms viz, SO_4_^2-^, SO_3_^2-^ and H_2_S through reductive pathway utilizing ATP Sulfurylase (APS), APS Reductase (APR) and Sulfite Reductase (Brychkova et al., 2013; Gonzalez Gordo et al., 2020). An enzyme Sulfite Oxidase is responsible for regeneration of sulfate from sulfite. The S from H_2_S now enter to the final process of assimilation into amino acid, and involve participation of OASTL (O-acetylserine(thiol)lyase), which produces cysteine (first sulfur-containing organic molecule generated by plants) after utilizing O-acetyl serine and sulfide, and produces acetate as byproduct. Liu et al., (2019) has characterized 8 OASTL in tomato, which were reported to localize at different sub-cellular compartments. However, cysteine can also revert to H_2_S by the participation of NIFS/NFS like protein (chloroplast and mitochondria), L/D-cysteine desulfurylase and Bifunctional L-3-cyanoalanine synthase/cysteine synthase D1 and D2, Bifunctional cystathionine gamma-lyase/ cysteine synthase and pyridoxal-5′ -phosphate dependent enzyme family protein, as reported in Arabidopsis (Gonzalez Gordo et al., 2020). The working model for the representation of diverse enzymes involved in H_2_S metabolism is depicted in Figure 1.

### 4.1. H_2_S metabolism in tomato utilizes different compartments

Our investigation suggested the participation of 29 proteins in tomato H_2_S metabolism (Table S1). The cytosolic H_2_S metabolism utilizes the activity of L-Cysteine desulfhydrase (SlLCD1 and SlLCD2), Cysteine synthase / O-acetyl-l-serine (thiol) lyase (SlOAS1-4, SlOAS6 and SlOAS9), and Selenium binding protein (SlSBP1). In the prediction, we could not find any sulfite reductase, and cystathionine gamma-synthase and cystathionine beta-lyase activity in the cytosol. The SBP is represented by only one member in tomato, contrary to three SBP in Arabidopsis (Gonzalez-Gordo et al., 2020). Liu et al., (2019) have also reported the localization of SlOAS4 in the cytoplasm; however, the location given for other cysteine synthase was somewhat different viz, SlOAS2 in membrane, SlOAS6 in nucleus and SlOAS9 in peroxisome. Interestingly, the clustering of genes encoding for SlOAS was observed at chromosome 1, namely *SlOAS1, SlOAS2, SlOAS3* and *SlOAS4*. The occurrence of SlLCD1 in the nucleus was also reported (Hu et al., 2020), which suggested that possibly the enzymes associated with H_2_S/cysteine metabolism are present in the nucleus by the activity of SlLCD1 (Hu et al., 2020) and SlOAS6 (Liu et al., 2020). But as we have also observed their presence in the cytosol through consensus score developed after analysis of 10 prediction tools, their activity in the cytosol (40% confidence) cannot be denied, though need further investigations. For SlOAS9, none of the prediction tool has demonstrated its presence in peroxisome. Further, we cannot deny the sub-cellular location of SlOAS9 in peroxisome, as this sub-cellular compartment is known for H_2_S metabolism. The peroxisome is the single membrane bound sub-cellular body actively engaged in nitro-oxidative metabolism (Corpas et al., 2019; Corpas and Palma, 2020). In tomato, no enzymatic source for H_2_S metabolism was observed in peroxisome, which is similar in Arabidopsis (Gonzalez-Gordo et al., 2020). In our prediction, we have observed the extracellular location of mercaptopyruvate sulfurtransferase (SlMST3) with high consensus score (40%), therefore, this protein has not been assigned to any sub-cellular location. Additionally, the location of Solyc04g055230.2.1 (SlCBL1) is also dicey, as it demonstrated opportunity to be localized equally, in almost all locations. Both SlMST3 and SlCBL1 are not considered for current model. Further, SlOAS3 was also not considered for current model (Fig. 1), as it does not produce functional OAS like protein, as reported previously (Liu et al., 2019).

The chloroplast is the site of sulfur assimilation into amino acids, which involves sulfate reduction by the activity of two enzymes (adenosine-5′ -phosphosulfate, APS and adenosine-5′-reductase, APR) into sulfite and to sulfide (by the activity of sulfite reductase) and cysteine production. The later includes the participation of cysteine synthase (OAS-(thiol)lyase, OAS-TL/ OASB) activity which combine H_2_S with O-acetylserine (OAS) to form cysteine (Birke et al., 2015). In our investigation, we have observed the participation of 12 enzymes as mentioned in section 3.1 (Table 2). Our investigation corroborated the findings of Liu et al., (2019), where they have demonstrated the location of SlOAS5 in chloroplast. Further, the SiR was also reported earlier to be localized in chloroplast (Brychkova et al., 2012). Except lipoyl synthase, all other enzymes are directly involved in the conversion of inorganic sulfur (sulfate and sulfite) to organic forms (Cysteine and homocysteine). The chloroplast H_2_S metabolism was observed similar in Arabidopsis and tomato; however tomato exhibited utilization of D-Cysteine desulfhydrase (as the activity demonstrated by closest ortholog in Arabidopsis, table 2), contrary to L-Cysteine desulfhydrase as reported in Arabidopsis (Gonzalez-Gordo et al., 2020).

The mitochondrial H_2_S metabolism play important role in mitigation of CN inducing toxicity due to the activity of mitochondrial bifunctional L-3-cyanoalanine synthase/cysteine synthase C1 (CYSC1). The enzyme was reported in Arabidopsis to perform detoxification of cyanide and involve conversion of cyanide and cysteine to β-cyanoalanine and hydrogen sulfide (Yamaguchi et al., 2000). In tomato, 5 enzymes were identified to be localized in mitochondria (section 3.1. Table 3). Though our analysis predicted the localization of SlMST4 with high confidence in chloroplast (60% Consensus score), but this enzyme may also be of significance in mitochondria (40% Consensus score). We here propose the dual localization of this protein in both chloroplast and mitochondria (Fig. 1). Though, we could not get the Cystathionine-β-lyase activity in either cytosol or mitochondria, but it is possible that SlCBL1 might function in these locations. Further, the SlOAS8 was functionally assigned as L-3-cyanoalanine synthase, which was showing closest homology with At3g61440 (AtCysC1) and suggested for this function by Liu et al., 2019. The experimental data also suggested the localization of SlOAS8 in mitochondria (Liu et al. 2019), thereby testify our prediction. Our analysis suggested that the mitochondrial H_2_S metabolism involves all the proteins as associated in Arabidopsis (Gonzalez-Gordo et al., 2020).

### 4.2. Proteins associated with H_2_S metabolism in tomato demonstrate compartment specific interaction

Protein–protein interactions (PPIs) signify to understand the important aspect of plant systems biology including metabolism. Further the elucidation of interaction networks provides critical insights into the regulatory aspects of plant proteins in plant developmental processes and its interactions with their environment (Struk et al., 2019). The interaction network of proteins associated with H_2_S metabolism was presented by Gonzalez-Gordo et al., (2020), which demonstrated the significant compartment specific interactome. The PPI analysis suggested the interaction of tomato proteins responsible for H_2_S metabolism in all the cellular compartments. Amongst, all proteins participated in the interaction network of tomato chloroplast; however, in cytosol except SlSBP1 all other proteins were found interacting. In cytosol, it was observed that no SlOAS interact with each other, where as these proteins exhibited interaction with both SlLCDs, which suggested the H_2_S/Cysteine metabolism related enzymes function in close association and exhibited regulation by interacting with each other viz, SlLCD1-SlOAS1, SlLCD1-SlOAS2, SlLCD1-SlOAS3, SlLCD1-SlOAS4, SlLCD1-SlOAS6, SlLCD1-SlOAS9, SlLCD2-SlOAS1, SlLCD2-SlOAS2, SlLCD2-SlOAS3, SlLCD2-SlOAS4, SlLCD2-SlOAS6 and SlLCD2-SlOAS9. In chloroplast, also the interaction was observed among enzymes responsible for H_2_S/Cysteine metabolism (SlDCD2-SlOAS5). Along, the other enzymes responsible for H_2_S metabolism also demonstrated considerable interactions. The SlLIP proteins of chloroplast also exhibited strong interaction with each other. Further, in mitochondria, two interacting groups were observed viz, SlOAS7-SlOAS8 and SlLIP2-SlNFS1-SlDCD1. Interestingly, no interaction of enzymes responsible for H_2_S/Cysteine metabolism was observed in mitochondria. In cystein biosynthesis, the interaction of OAS and SAT (Serine acetyl transferase), thus forming cysteine synthase complex as reported in Arabidopsis (Francois et al., 2006). Similarly, such interaction is also reported in tomato (Liu et al., 2019). Our finding offers the formation of possible complex (comprised of H_2_S production and consumption activty) for H_2_S/Cysteine metabolism, present in tomato cell. The study signifies the possibility about the highly coordinated network of proteins associated with H_2_S metabolism in tomato, which is responsible for sulfur metabolism including H_2_S/cysteine homeostasis.

### 4.3. Genes associated with H_2_S metabolism exhibited modulation under development and biotic stress

The genes encoding enzymes responsible for either H_2_S biosynthesis or consumption were utilized for the expression analysis. Over all the assessment of cytosolic (*SlLCD1, SlOAS1, SlOAS2, SlOAS3, SlOAS4, SlLCD2, SlOAS6* and *SlOAS9*), chloroplastic (*SlDCD2, SlNFS2, SlOAS5* and *SlSiR1*) and mitochondrial (*SlDCD1, SlOAS7* and *SlNFS1*) genes were assessed. In development stages, the *SlLCD1/SlOAS6, SlOAS5* and *SlNFS1* were observed as highly expressed genes in cytosol, chloroplast and mitochondria, respectively; however, they exhibited peak expression in different tissues as mentioned in results section (Fig. 3). The high activity of H_2_S biosynthetic genes *SlLCD1* and *SlLCD2* during stages of fruit development is possibly compensated by activity of cysteine synthase activity (mostly offered by *SlOAS6*). The findings corroborated with the results of Hu et al., (2020), which exhibited accelerated fruit ripening in *SlLCD1* silenced and CRISPR/Cas9 mediated SlLCD1 gene-edited mutant. The study has also reported the high *SlLCD1* expression during fruit ripening and suggested the role of *SlLCD1* and H_2_S in the regulation of fruit ripening, which is similar to our study. The increase in cytosolic H_2_S and L-cysteine desulfhydrase activity was also observed in sweet pepper (*Capsicum annuum* L.) fruit ripening (Muñoz-Vargas et al., 2018). Further, the substantial production of cytosolic *SlLCD1* and *SlOAS6* in all tissues, demonstrated that the encoded proteins of these genes possibly play crucial role and mainly responsible for the H_2_S mediated response in plant development.

During biotic stress, the significant modulation in the expression of H_2_S production and consumption associated genes was also observed. Interestingly, *SlLCD1, SlLCD2, SlOAS9, SlSiR1* and *SlNFS1* were observed as late responsive and their up-regulation was mostly observed at 6 h of PAMP treatment/bacterial infection, which suggested the significance of H_2_S during plant-pathogen interaction. Contrary to this, *SlOAS7* was observed as early responsive. Vojtovic et al., (2021) have reviewed the contribution of H_2_S in plant defense. A study of knockout mutants to plant pathogen in Arabidopsis demonstrated that the high resistance to biotrophic and necrotrophic pathogens in *des1* mutant and sensitivity in *oas-a1* mutant (Alvarez et al., 2012). In our study, the cytosolic *SlOAS6* and *SlOAS9* expression was down-regulated during early stage of PAMP treatment, which otherwise up-regulated at later stage, which might be related with the contribution of encoded cysteine synthase activity in PAMP resistance at 6 h, which was most prominent in case of csp22 and flgII-28, respectively. However, the *SlLCD1* though demonstrated similar trend, but exhibited up-regulation with few PAMP such as LPS. Further, in chloroplast and mitochondria, the contribution of *SlOAS5-SlNFS2* and *SlOAS7-SlNFS1* play a significant role in H_2_S consumption, and production respectively (Fig 3). In response to bacterial infection, up-regulation of *SlOAS2, SlOAS6* and *SlOAS9* demonstrated up-regulation; however, *SlOAS7* showed down-regulation for *Pseudomonas* sps, and up-regulation for *A. tumifaciens* at 6 h. Among gene responsible for H_2_S production, *SlNFS2* demonstrated down-regulation in response to *P. fluorescence* and *P. putida; SlNFS1* demonstrated up-regulation mainly for *Pseudomonas* sps and *SlDCD1* exhibited down-regulation with *P. syringae* and *P. putida* at 6 h duration. The finding clearly demonstrates the significant modulation of genes associated with H_2_S production and consumption activity and thus testifies their contribution in related events.

## 5. Conclusion

In nut shell, the exploration of genes associated with H_2_S metabolism was explored in tomato. Over all 29 proteins were identified, which are related to tomato H_2_S metabolism. The study also offers an insight on their potential interaction at sub-cellular level. The results demonstrated the significant interaction among the proteins present in cellular compartments such as cytosol, chloroplast and mitochondria. Further, the expression of selective genes was studied under diverse developmental and biotic stress conditions. The specific expression pattern of genes associated with H_2_S production and consumption demonstrated their occurrence in critical events related to development, particularly fruit development. Moreover, their regulation under biotic stress (PAMP and bacterial treatment) offers considerable support to visualize and explore their function in relation to plant-pathogen interaction. Over all, we believe that investigation of this important metabolism in economically important horticulture crop (tomato) will open new prospect for the investigation of various biological functions.

## Author contributions

VS^1^: Framing the concept; AAC^1^, SM^2^, VS^3^ and VS^1^: Performed analysis and Manuscript writing; AAC^1^, SM^2^ and VS^1^: Figures and tables preparation; VS^1^ and VS^3^: Correction and edited the manuscript.

## Acknowledgements

AAC^1^ and VS^1^ acknowledge Central University of Jammu for providing working facility. VS^1^ also acknowledges UGC, India for UGC-BSR Research Start-up grant.

## Figure and Table Legends

Table S1 The sub-cellular localization prediction of tomato proteins related to H_2_S metabolism.

