## Supplementary material for "Elucidation of sub-cellular H_2_S metabolism in *Solanum lycopersicum* L. and its assessment under development and biotic stress": Table S1

Table S1: The sub-cellular localization prediction of tomato proteins related to H_2_S metabolism.

| **Protein ID**  **(Solgenomics)** | **Wolf P SORT** | **Deeploc 1.0** | **Chloro .p** | **TargetP 2.0** | **Plant-mSubP** | **BUSCA** | **LOC TREE 3** | **DeepMito** | **LOCALIZER** | **CELLO2GO** | **Sub-cellular Location (% Confidence)** |
| --- | --- | --- | --- | --- | --- | --- | --- | --- | --- | --- | --- |
| Solyc01g008900.4.1 | Nucleus | Mitochondrion | - | Other | Celmemb | Nucleus | Mitochondrial outer membrane | No | Chloroplast , nucleus | Mitochondrial | Mitochondria (30)  Nucleus (30) |
| Solyc01g068160.4.1 | Cytoplasmic | Cytoplasm | - | Other | Cytoplasm | Cytoplasm | Mitochondrion | No | Nucleus | Cytoplasm | Cytoplasm(50) |
| Solyc01g094790.3.1 | Chloroplast | Mitochondrion | - | Mitochondrial transfer peptide | Mitochondrion | Mitochondrion | Mitochondrion | No | Mitochondrion | Mitochondrial | Mitochondria (70) |
| Solyc01g097920.3.1 | Cytoplasmic | Peroxisomes | - | Other | Cytoplasm | Cytoplasm | Mitochondrion | No | - | Cytoplasm | Cytoplasm(40) |
| Solyc01g097930.3.1 | Cytoplasmic | Cytoplasm | - | Other | Cytoplasm | Cytoplasm | Cytoplasm | No | - | Cytoplasm | Cytoplasm(60) |
| Solyc01g097940.1.1 | Extracellular | Cytoplasm | - | Other | Plastid | Cytoplasm | Cytoplasm | No | - | Chloroplast | Cytoplasm(30) |
| Solyc01g097950.3.1 | Cytoplasmic | Cytoplasm | - | Other | Plastid | Cytoplasm | Cytoplasm | No | - | Cytoplasm | Cytoplasm(50) |
| Solyc02g067180.3.1 | Chloroplast | Plastid, soluble | - | Chloroplast transfer peptide | Plastid | Mitochondrion | Chloroplast | Mitochondrial | Chloroplast | Chloroplast | Chloroplast(70) |
| Solyc02g078990.4.1 | Vacuolar | Plastid | cTP | Other | Extracellular | Cytoplasm | Mitochondrion | No | Nucleus | Cytoplasm | Cytoplasm(20)  Chloroplast(20) |
| Solyc02g091900.4.1 | Chloroplast | Mitochondrion | - | Mitochondrial transfer peptide | Mitochondrion | Mitochondrion | Mitochondrion | Mitochondrial | Nucleus | Mitochondrial | Mitochondria (70) |
| Solyc03g098230.3.1 | Cytoplasmic | Plastid | CTP | Chloroplast transfer peptide | Plastid | Chloroplast membrane | Mitochondrion | Mitochondrial | Chloroplast , nucleus | Chloroplast | Chloroplast(70)  Mitochondria (20) |
| Solyc04g055230.2.1 | Cytoplasmic | Mitochondrion | - | Other | Celmemb | Nucleus | Chloroplast | No | - | Extracellular | Cytoplasm(10)  Mitochondria (10)  Chloroplast(10)  Extracellular(10)  Nucleus (10)  Cel memb (10) |
| Solyc04g058120.3.1 | Nucleus | Mitochondrion | - | Other | Mitochondrion | Chloroplast | Mitochondrion | No | Mitochondrion | Mitochondrial | Mitochondria (50) |
| Solyc05g007590.4.1 | Cytoplasmic | Mitochondrion | - | Other | Cytoplasm | Cytoplasm | Chloroplast | No | - | Cytoplasm | Cytoplasm(40) |
| Solyc05g055000.4.1 | Chloroplast | Plastid | CTP | Chloroplast transfer peptide | Mitochondrion | Mitochondrion | Chloroplast | Mitochondrial | Chloroplast | Mitochondrial | Chloroplast(60)  Mitochondria (40) |
| Solyc06g009850.3.1 | Chloroplast | Plastid | CTP | Other | Golgi | Cytoplasm | Mitochondrion | Mitochondrial | Nucleus | Mitochondrial | Chloroplast(30)  Mitochondria (30) |
| Solyc06g009860.1.1 | Extracellular | Extracellular | - | Other | Celmemb | Extracellular space | Mitochondrion | No | - | Extracellular | Extracellular(40) |
| Solyc07g054540.4.1 | Nucleus | Plastid | CTP | Other | Plastid | Chloroplast | Mitochondrion | Mitochondrial | Chloroplast , nucleus | Nucleus | Chloroplast(50) |
| Solyc07g065470.4.1 | Cytoplasmic | Cytoplasm | - | Other | cytoplasm | Cytoplasm | Chloroplast | No | - | Chloroplast | Cytoplasm(40) |
| Solyc07g066580.3.1 | Chloroplast | Mitochondrion | cTP | Chloroplast transfer peptide | Plastid | Chloroplast membrane | Mitochondrion | Mitochondrial | Mitochondrion | Chloroplast | Chloroplast(60)  Mitochondria (40) |
| Solyc08g014340.3.1 | Chloroplast | Plastid | cTP | Chloroplast transfer peptide | Plastid | Chloroplast membrane | Chloroplast | Mitochondrial | Chloroplast | Chloroplast | Chloroplast(90) |
| Solyc08g066620.3.1 | Chloroplast | Cytoplasm | - | Other | Endoplasm | Chloroplast | Chloroplast | Mitochondrial | Mitochondrion | Plasmamembrane | Chloroplast(30)  Mitochondria (20) |
| Solyc08g068440.4.1 | Extracellular | Mitochondrion | - | Other | Mitochondrion | Chloroplast | Mitochondrion | No | Chloroplast | Chloroplast | Mitochondria (30)  Chloroplast(30) |
| Solyc09g082060.3.1 | Cytoplasmic | Cytoplasm | - | Other | Cytoplasm | Cytoplasm | Cytoplasm | No | - | Cytoplasm | Cytoplasm(60) |
| Solyc09g092430.3.1 | Cytoplasmic | Cytoplasm | - | Other | Cytoplasm | Nucleus | Cytoplasm | No | - | cytoplasm | Cytoplasm(50) |
| Solyc10g012370.3.1 | Mitochondrial | Mitochondrion | - | Mitochondrial transfer peptide | Mitochondrion | Mitochondrion | Mitochondrion | Mitochondrial | Mitochondrion | Mitochondrialchloroplast | Mitochondria (90) |
| Solyc10g079720.2.1 | Chloroplast | Plastid soluble | CTP | Chloroplast transfer peptide | Plastid | Chloroplast | Chloroplast | Mitochondrial | Chloroplast | Chloroplast | Chloroplast (90) |
| Solyc11g065620.2.1 | Cytoplasmic | Plastid | CTP | Chloroplast transfer peptide | Plastid | Mitochondrion | Chloroplast | Mitochondrial | Chloroplast , nucleus | Mitochondrial | Chloroplast(60)  Mitochondria (30) |
| Solyc12g099700.3.1 | Chloroplast | Plastid | - | Other | Plastid | Mitochondrion | Chloroplast | Mitochondrial | - | Chloroplast | Chloroplast(50) |
